# A structural equation model for imaging genetics using spatial transcriptomics

**DOI:** 10.1101/253443

**Authors:** Sjoerd M.H. Huisman, Ahmed Mahfouz, Nematollah K. Batmanghelich, Boudewijn P.F. Lelieveldt, Marcel J.T. Reinders

## Abstract

Alzheimer’s disease is a neurodegenerative disorder that causes changes in the structure of the brain, observable with MRI scans, and that has a strong heritable component, reflected in the DNA. Imaging genetics deals with such relationships between genetic variation and imaging variables, often in a disease context. The complex relationships between brain volumes and genetic variants have been explored both with dimension reduction methods and model based approaches. However, these models usually do not make use of the extensive knowledge of the spatio-anatomical patterns of gene activity. We present a method for integrating genetic markers (single nucleotide polymorphisms) and imaging features, which is based on a causal model and, at the same time, uses the power of dimension reduction. We use structural equation models to find latent variables that explain brain volume changes in a disease context, and which are in turn affected by genetic variants. We make use of publicly available spatial transcriptome data from the Allen Human Brain Atlas to specify the model structure, which reduces noise and improves interpretability. The model is tested in a simulation setting, and applied on a case study of the Alzheimer’s Disease Neuroimaging Initiative.

## Introduction

The aim of imaging genetics studies is to find associations between genetic variants and imaging features, often in a disease context [1]. This scheme extends beyond traditional genome wide association studies (GWAS) by identifying genetic associations of imaging biomarkers with the assumption that these biomarkers are a more direct reflection of the genetic effects. Thus, they could provide a stronger association signal [2]. Additionally, the identified associations are likely to provide new insights into the underlying disease mechanisms as well as new hypotheses about the anatomical and/or functional locations involved in complex diseases [3].

So far, imaging genetics studies have been largely focused on the brain [1,3–6], despite efforts to extend their application to other fields [7]. Several large consortia have gathered data from thousands of subjects to understand the effects of genetic variants on brain structure and function [8]. One of the hallmark sources for imaging genetics studies is the Alzheimer’s Disease Neuroimaging Initiative (ADNI) database [9]. This database contains single nucleotide polymorphism (SNP) and structural MRI data for Alzheimer’s patients, individuals with late mild cognitive impairment, and cognitive normal controls.

One of the largest challenges facing imaging genetics studies is the statistical power needed to identify reliable associations. In a typical GWAS, researchers have to correct for the number of independent tests performed (i.e. number of independent SNPs tested) in order to limit the number of false positive discoveries. However, a genome-wide brain-wide imaging genetic study will not only have to correct for the number of independent SNPs, but also for the number of independent imaging features tested. This requirement yields most of the studies underpowered to identify reliable associations. One of the largest imaging genetics studies [10] analyzed over 30,000 individuals within the Enhancing Neuro Imaging Genetics through Meta-Analysis (ENIGMA) consortium. They performed a genome wide association of SNPs with seven brain volumes, and identified only eight genome wide significant SNPs.

Despite the high dimensionality of the imaging data (millions of voxels), the actual number of independent tests for which we need to correct in an imaging genetics study is far smaller than the number of voxels. Due to the spatial relationships between voxels, measurements from neighboring voxels are usually highly correlated. A common approach is to test genetic associations for anatomically defined brain regions [2]. Several studies have shown that both neuroanatomical parcellation and connectivity of the brain are strongly reflected in gene expression patterns across the brain [11–13]. The public availability of brain transcriptome atlases from the Allen Institute for Brain Science [14] provides an opportunity to use these transcriptional signatures to group brain regions, limiting the number of effective tests.

Several methods have been proposed to identify associations between genetic variants and imaging features by applying dimension reduction, such as variations of canonical correlation analysis [15], and independent component analysis (commonly used in a functional MRI context) [4]. Others have opted to model the interactions between the different data types explicitly. Both [16] and [17] pose graphical Bayesian models which capture a more mechanistic causal view of the data. These models consist of a directed acyclic graph, which can easily be made to incorporate covariates, including possible confounding factors. Both studies use relatively small candidate SNP sets, because they aim for understanding SNP brain relationships rather than the discovery of genome wide associations. However, these Bayesian models are quite challenging to specify and fit.

In this work, we propose a method to identify associations between candidate genetic variants and imaging features allowing for the incorporation of prior knowledge. The proposed method combines a graphical model with dimension reduction to model the effect of SNPs on brain imaging features through a set of latent variables. We use a maximum likelihood structural equation modelling (SEM) approach to find the edge weights of our model [18]. By performing dimensionality reduction within the model, we reduce the number of parameters to be estimated. In addition, the model allows for easy incorporation of information from the Allen Human Brain Atlas [12] to inform the grouping of brain regions based on the similarity of their transcriptional profiles.

## Materials and methods

The interplay between genetic variation, brain anatomy, and disease symptoms is complex. We use a structural equation model with latent variables [18] to model these relationships. We pose that the genetic variation is exogenous, in other words: the genetic variation in a study population is not caused by disease or brain anatomy. This variation does have an effect on the brain. For example, in Alzheimer’s disease, genetic variants may influence the immune response and amyloid *β* concentrations in the brain, which may in turn lead to shrinkage in several brain areas [19]. Large scale imaging initiatives, such as ADNI, offer a possibility to study this shrinkage of brain regions. This can be estimated from MRI data of diseased individuals and controls, and expressed in cortical thickness and subcortical volume measurements.

In our graphical model, we define groups of brain regions, based on the transcriptional profiles of these areas in the healthy brain. Areas that share patterns of gene expression in a normal brain may be similarly affected by genetic variations. For each of the region groups, we introduce one latent variable. This latent variable is affected by the genetic variations, and causes changes in relevant brain regions. This makes our model similar to principal component analysis (PCA) on sets of brain regions, combined with a regression for the latent variables.

### Variables used

We model the relationship between single nucleotide polymorphisms (SNPs) and brain region measurements. Let g_*i*_ ∈ℝ^*p*^ be a vector of centred (zero-mean) SNP values, and x_*i*_ ∈ℝ^*q*^ a vector of centred (zero-mean) and scaled (sd = 1) brain region measurements, both for individual *i*. The reason both types of measurements are centred, is to eliminate intercepts from the model. The brain measurements are, in addition, scaled to unit variance to compensate for the considerably larger variance in thickness or volume for larger brain areas. The genetic variants and brain measurements are connected in the model by a set of latent variables, z*_i_*∈ℝ*^m^*.

In addition to the variables included in the model, we have two other sources of information. In defining the model structure, we make use of external information on the brain region measurements, in the form of brain region groups with a shared transcriptional profile. These brain regions can be defined based on spatial gene expression data of the healthy adult brain. Finally, the goal is to understand disease related phenotypes. The disease labels are not used in the modelling stage. However, we hypothesize that if the variation in the data is related to a disease state, the latent variables will reflect this. After model fitting, we therefore associate each individual’s estimated latent variable score to his or her disease status.

### The graphical model

We model the relationship between brain SNP values and brain region measurements in a structural equation model (SEM). It consists of two parts. The first part is a linear model for brain region measurements as a function of the latent variables,

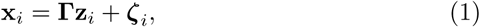

where x*_i_* contains the observed brain region measurements, z*_i_* the latent variables, and ζ*_i_* is a zero-mean normally distributed error variable. The matrix **⌈** contains the weights of the latent variables that explain the brain region measurements. The second part of the SEM is a linear model for these latent variables as a function of the SNP values,

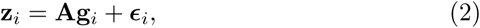

where **g***_i_* contains the observed SNP measurements, and *∈_i_* is a zero-mean normally distributed error variable. The matrix **A** contains regression weights, representing the effects of the SNPs on the latent variables. Combined, these equations mean that region changes are viewed as a manifestation of the latent values, while the SNP values are considered causal to them. The latent variables represent some intermediate phenotype, related to the molecular state of the connected brain regions.

The number of latent variables is equal to the number of brain region groups, which are defined based on external spatial gene expression data. A region group contains the brain regions with a similar transcriptional profile, as these may react similarly to differences in genetic background. We restrict each latent variable to only predict the brain region measurements for its own region group. This results in a restriction on the weight matrix **Γ**, where each latent variable (corresponding to a column in **Γ**) has a unique set of non-zero entries. Fig 1 shows the model for two latent variables, where we can see that each latent variable is connected to its own set of brain regions.

**Fig 1.**
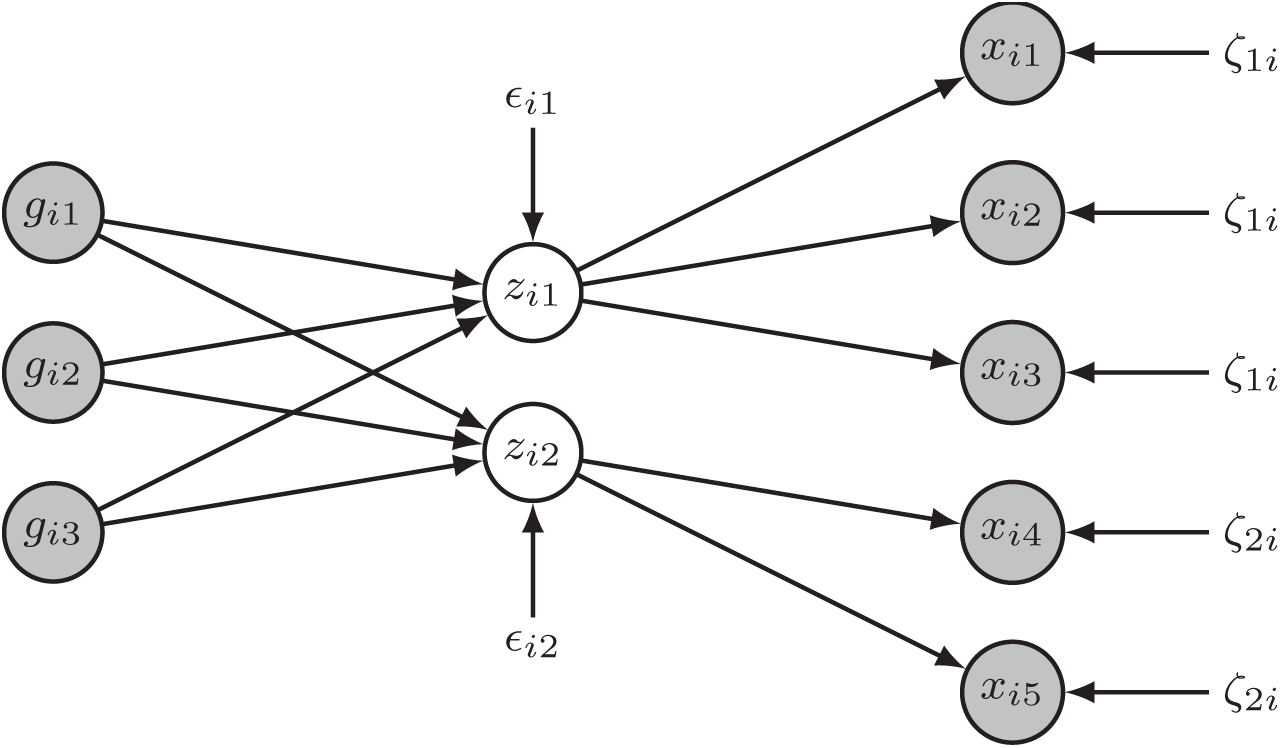
The graphical structural equation model. Observed variables are shown in grey circles, latent variables in white circles, and error variables without circles. This example contains two latent variables, both with their own set of observed brain region measurements. This structure, where the latent variables define groups of region measurements, is defined by prior knowledge on these brain regions. We use spatially resolved gene expression data of the healthy human brain to define these region groups.

### Model implied covariance

In linear Gaussian structural equation modelling, we learn the parameters of a model by optimising the correspondence between the observed covariance **S** (from the data), and the model implied variance **Σ**. The model implied variance can be divided in a block matrix, by defining

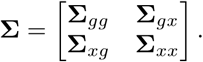

Note that this implied covariance does not contain any components for the latent variables in **z**. The latent variables are not observed, and therefore we can not use their observed covariance in fitting the model.

The elements of the implied covariance can be parametrised in terms of the model coefficients. The first element is

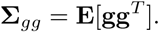

This is the covariance of the SNPs (since these values are centred). The SNPs are exogenous in our model: g does not have any causal variables within our model. As a result, the implied covariance of the SNPs is not parametrised in terms of model coefficients. We can estimate this covariance term simply by taking the observed covariance between the SNPs.

The next element of the implied covariance matrix is

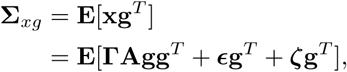

and similarly

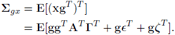

The final element of the implied covariance matrix is the model implied covariance among the brain regions. This is given by

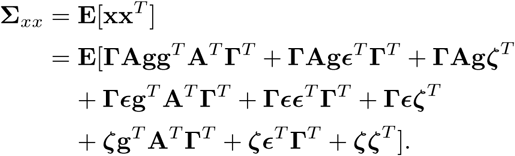

### Model assumptions and estimation

Some elements of the implied covariance are often assumed to be zero. These assumptions lead to a strong simplification of the implied covariance. It is common in a regression setting to pose that the predictor variables and error variables are independent. In our case, the error independence assumption leads to g*∈^T^* = ∈g*^T^* = 0. In addition, we assume that the errors in the brain region predictions (equation (1)) are independent of the errors in the latent variable predictions (equation (2)). This means that ζ*∈^T^* = ∈ζ*^T^* = 0. Finally, we assume that the errors in brain region prediction are independent of the SNPs, so gζ*^T^* = ζg*^T^* = 0.

As a result of these assumptions, the full implied covariance matrix of the model reduces to

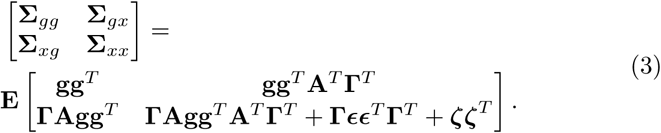

For normally distributed data, the maximum likelihood estimate of the covariance matrix is

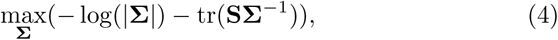

where is **S** the observed covariance matrix. The SNP data we use is discrete, and can therefore not be considered normally distributed. To compensate for this, we will estimate robust standard errors. In equation (4), the covariance **Σ** is parametrised according to equation (3), so we can perform the optimisation over the parameter values.

Model fitting is performed in the *lavaan* package in *R* [20]. For identifiability, we fix the loading of the first brain region measurement per region group (latent variable) to 1. This does not only fix the scales of the latent variables, but it also has the advantage that the resulting latent variables will have the same direction of effect as the first brain region measurement. For example, a reduction in volume of the first brain region will result in a reduction in the corresponding latent variable. All the error variances on the brain region measurements (variance of ζ) are assumed to be equal, which is the same as in principal component analysis.

The model fit in *lavaan* yields estimates for **Γ**, **A**, and the covariance matrices of the error variables **∈** and ζ. Each of these parameter estimates is provided with robust p-values (for the hypothesis of being equal to zero), when using the *MLM* estimation procedure [20]. Using the estimated model parameters, one can then calculate unbiased Bartlett scores for the latent variables [21].

### Data

#### Simulated data

The model is evaluated both on simulated and real data. In the simulation, we first generated SNP values (g*_i_*) in accordance with Hardy-Weinberg equilibrium. The minor allele frequencies were independently drawn from a beta distribution with shape parameters *α* = 1 and *β* = 2. Then we simulated latent variables (z_i_) as a linear combination of the SNP values, with Gaussian noise (*sd* = 2). Each of these latent variables determined the region measurements (x*_i_*) of a set of regions (a region group), with added Gaussian noise (*sd* = 2). This part of the simulation is in line with equations (1) and (2) and Fig 1. Finally, we used a logistic model in which a linear combination of some of the latent variables determined the probability of observing a phenotype. These binary phenotypes (disease versus healthy) were then drawn from a Bernoulli distribution.

We simulated 100 independent datasets for 500 individuals. Each time, we set the number of SNPs to 20 and the number of latent variables (and therefore region groups) to 5. We randomly selected 10 SNP-to-latent weights (**A**) to be either 1 or −1. The 5 region groups contain 20, 10, 10, 5, and 5 regions respectively, for a total of 50 brain region measurements. Each latent variable has latent-to-brain-region weights (in **Γ**), which were uniformly sampled between 0.5 and 1.5. All other elements of **Γ** were set to zero, which effectively restricts each latent variable to affect only its own region group. Finally, two out of the five latent variables were randomly selected to affect the disease probability, with weights of either 10 or −10. All other latent-to-phenotype weights were set to zero.

### ADNI data and preprocessing

The real data used in the preparation of this article were obtained from the Alzheimer’s Disease Neuroimaging Initiative (ADNI) database (adni.loni.usc.edu)[9]. The ADNI was launched in 2003 as a public-private partnership, led by Principal Investigator Michael W. Weiner, MD. The primary goal of ADNI has been to test whether serial magnetic resonance imaging (MRI), positron emission tomography (PET), other biological markers, and clinical and neuropsychological assessment can be combined to measure the progression of mild cognitive impairment (MCI) and early Alzheimer’s disease (AD). For up-to-date information, see www.adni-info.org.

The ADNI database contains measurements on a large number of cognitive normal (CN) controls, individuals with late mild cognitive impairment (LMCI), and individuals with Alzheimer’s disease (AD). The measurements in the database include patient demographics, raw and processed MRI data, biomarker data and SNP data. For the brain volumes we made use of the UCSF cross-sectional FreeSurfer (Version 4.3) cortical thickness and white matter parcellation measurements. For the SNPs we made use of the ADNI 1 Illumina Human 610-Quad BeadChip data, with imputation as previously described [17]. In the end, we selected volumes, SNPs and diagnoses for 746 individuals. This data was split in two equal parts of 373 individuals, one as a training and one as a validation, to prevent over-fitting in the modelling process.

Our methodology is not suited to genome wide analysis. Instead, it tries to find the effects of specific SNPs on a set of latent variables. As candidate SNPs we selected a set of 35 polymorphisms associated with Alzheimer’s disease according to the International Genomics of Alzheimer’s Project (IGAP) study results [23]. IGAP is a two-stage GWAS on individuals of European ancestry for Alzheimer’s disease. In stage 1, IGAP used genotyped and imputed data on 7,055,881 SNPs of 17,008 Alzheimer’s disease cases and 37,154 controls. In stage 2, 11,632 SNPs were genotyped and tested for association in an independent set of 8,572 Alzheimer’s disease cases and 11,312 controls. Finally, a meta-analysis was performed combining results from stages 1&2. We selected the known SNPs, stage1 discoveries, and stage 1&2 discoveries from table 2, and the suggestive SNPs from supplementary table 4 of [23].

The volume data was present for 112 regions. We corrected it for individual age, gender, and whole brain volume (using linear regression),with the goal of maintaining all meaningful variation in brain region volumes, possible related to the disease phenotype. Out of the 112 regions, 105 regions were divided into nine region groups, see Table 1 and Supplemental Table 1. We defined these regions groups based on the transcriptional profiles in the healthy adult human brain, as provided by the Allen Atlas [12]. The division into region groups was made based on a non-linear dimension reduction of the gene expression values, and is shown in figure 2 of [24].

**Table 1.** The used region groups, with the regions they contain.

| Region group code | ABA region |
| --- | --- |
| CrCortex | cingulate gyrus |
| CrCortex | frontal lobe |
| CrCortex | insula |
| CrCortex | middle frontal gyrus |
| CrCortex | occipital lobe |
| CrCortex | parahippocampal gyrus |
| CrCortex | parietal lobe |
| CrCortex | temporal lobe |
| Hippocam | hippocampal formation |
| Amygdala | amygdala |
| Striatum | striatum |
| DorsThal | dorsal thalamus |
| SubCort1 | myelencephalon |
| SubCort2 | globus pallidus |
| SubCort2 | white matter |
| ClCortex | cerebellar cortex |
| SulcSpac | sulci & spaces |

## Results

### Simulation

To evaluate the performance of our model, the SEM was fitted to each of the simulated datasets. We considered two measures for model comparison. First, we set out to assess the prediction of phenotypes from the latent variables, with a logistic regression. In each of the 100 simulated datasets, we estimated the latent variable scores, and used only those to predict the phenotype. For each of these 100 models, we obtained an Akaike information criterion (AIC) value. We compared our model to several logistic regression models that use only the simulated data, instead of the SEM estimated latent scores. The first alternative model uses only all the brain region measurements, the second only all the SNP measurements, and the third a combination of all regions and SNP measurements. As a fourth alternative model, we performed a PCA on the volume measurements, and extracted the first five principal components. Fig 2 shows that, on average, our latent variables obtain a lower AIC than models using either all brain region data, all SNP data, or both. The model using the first five principal components of the brain region data is most similar to our model, and it only has a slightly lower AIC on average than our model.

**Fig 2.**
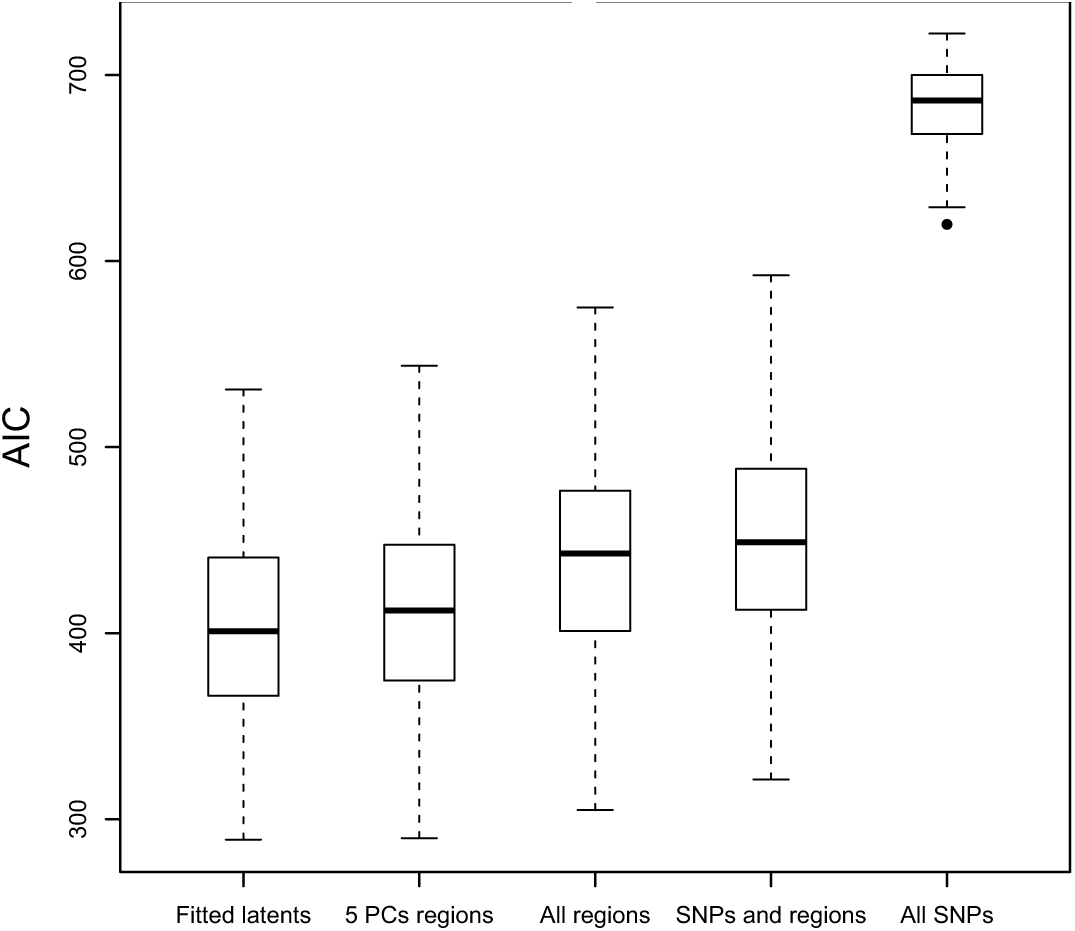
Simulation AIC model fit. The logistic regression models use either the SEM estimated latent variable scores (*Fitted latents*), the first five principal components of the brain region data (*5 PCs regions*), all brain region data, all SNP and brain region data, or all SNP data

The second measure for model comparison is the ability to retrieve the correct SNPs. In each of our simulation datasets, two of the five latent variables have an effect on the phenotype (disease status). All SNPs that affect either of these two latent variables effectively impact the phenotype. We consider those SNPs to be the SNPs with a true effect. We now consider how these SNPs are ranked for importance in our SEM analysis, and two alternative approaches. From our SEM fit, we extracted the robust SNP p-values for predicting the latent variables (so the p-values for the estimates in **A**). These give an impression of the importance of a SNP in predicting the latent variables. In addition, we used the latents’ logistic regression p-values for the phenotype. These show the importance of a latent variable in predicting the phenotype. As a result, the path from a SNP to the phenotype contains two p-values per latent variable: one for the latent variable prediction, and one for the phenotype prediction.

We considered combining these p-values in two ways: 1) for each SNP we took the maximum p-value of the two per latent variable, and then the minimum p-value over the five latent variables; or 2) for each SNP we used Fisher’s method [22] to combine the two p-values per latent variable (−2 Σ log(*p_i_*)), and then took the minimum p-value over the five latent variables. Note that Fisher’s method is meant for p-values testing the same null-hypothesis, which is not the case here. Both methods yield a score (p-value) for SNP importance. The order of these scores defines an ordering of the SNPs. Now, using the set of SNPs with a known true effect, we can construct a receiver operating characteristic curve for SNP retrieval, and the corresponding area under the curve (AUC).

We compared the performance of our methodology to a straightforward modelling approach: a logistic regression to predict the phenotype from the SNPs. This was performed both in a univariate way (as in a GWAS), and a multivariate way. Fig 3 shows the performance of our SEM based methods, using the maximum p-value per SNP-latent combination (*SEM max*) or using Fisher’s method (*SEM Fisher*), and of the GWAS-like approaches. The *SEM max* method has the highest average AUC, indicating that it is best able to rank the SNPs on their importance for the phenotype. Note that the *SEM Fisher* method has the disadvantage that either a strong SNP-to-latent or a strong latent-to-phenotype effect can lead to a low combined p-value, regardless of the other value. The observed difference between the univariate an multivariate approach is very small, which is to be expected since the simulated SNP values are independent.

**Fig 3.**
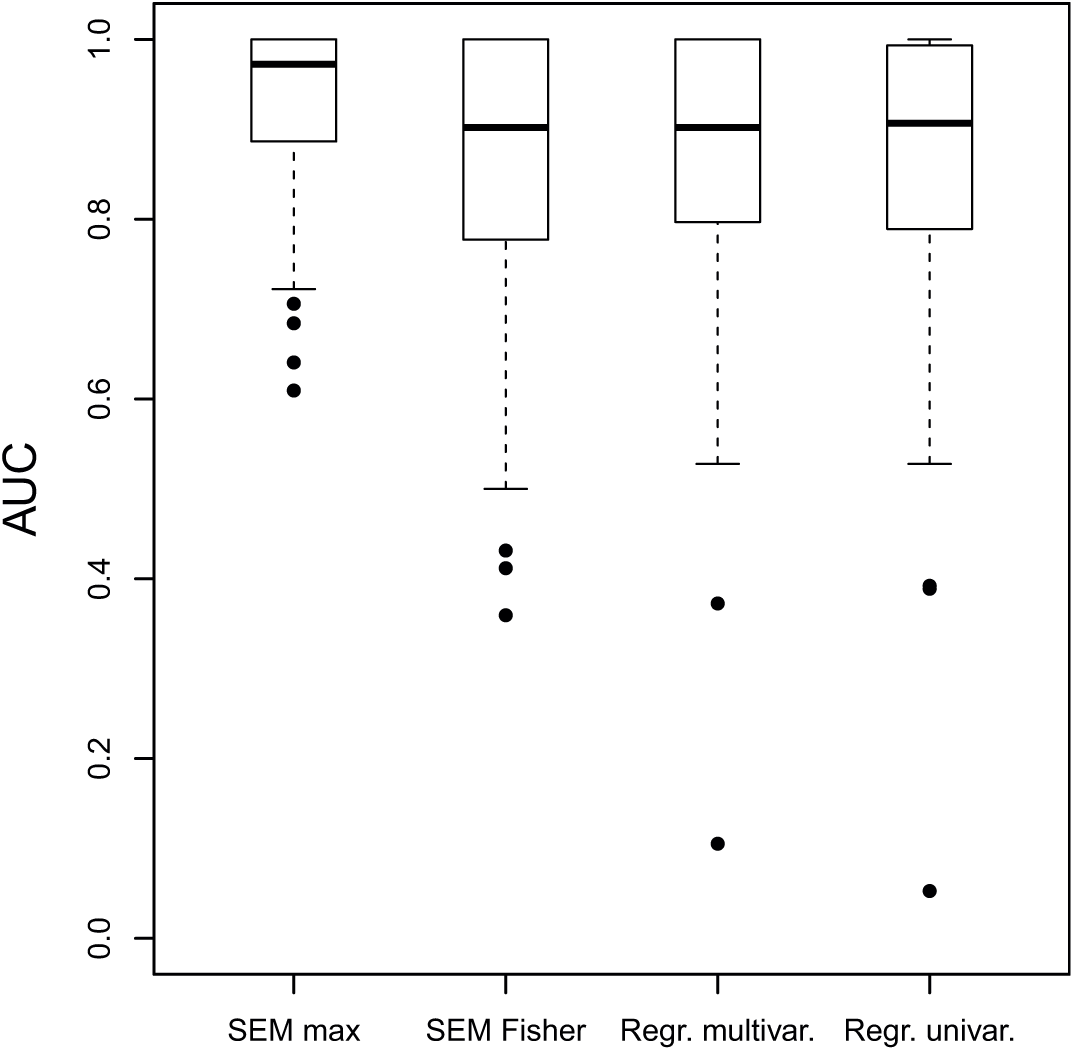
Simulation AUC for SNP selection. Shown are the results for two methods of p-value integration for our model (*SEM max* and *SEM Fisher*), for multivariate logistic regression, and univariate logistic regression. A high AUC means that the method correctly ranks the importance of the SNPs for the phenotype (disease state).

### ADNI application

We apply our methodology to the Alzheimer’s Disease Neuroimaging Initiative (ADNI) data [9]. We selected 35 SNPs and 105 brain region volumes for 746 individuals. The brain regions were divided into nine region groups based on the gene expression patterns of matching brain areas in the healthy human brain [12,24]. Each of the nine brain region groups has one corresponding latent variable, and each latent variable has a unique set of brain region measurements attached to it. Supplemental Fig 1 shows the volume loadings for each of the latent variables. Since the first loading for each latent variable is set to 1, the latent variables will have the same direction of effect as this variable. All but two of the region volumes have a positive loading. Two regions in the subcortical group 2 (*SubCort2*) are negatively correlated to the latent variable scores, reflecting a more heterogeneous signal in this group.

Fig 4 shows the association between the nine latent variables and the selected SNPs. Only those SNPs are shown that have a nominally significant (*p* < 0.05) association with at least one of the latent variables. The strongest effect is for rs429358, located in *APOE*, on the hippocampal region group (Bonferroni corrected *p* = 2.28 · 10^−4^). In the validation set, here used as a replicate, this effect was again significant (Bonferroni corrected *p* = 8.66 · 10^−3^).

**Fig 4.**
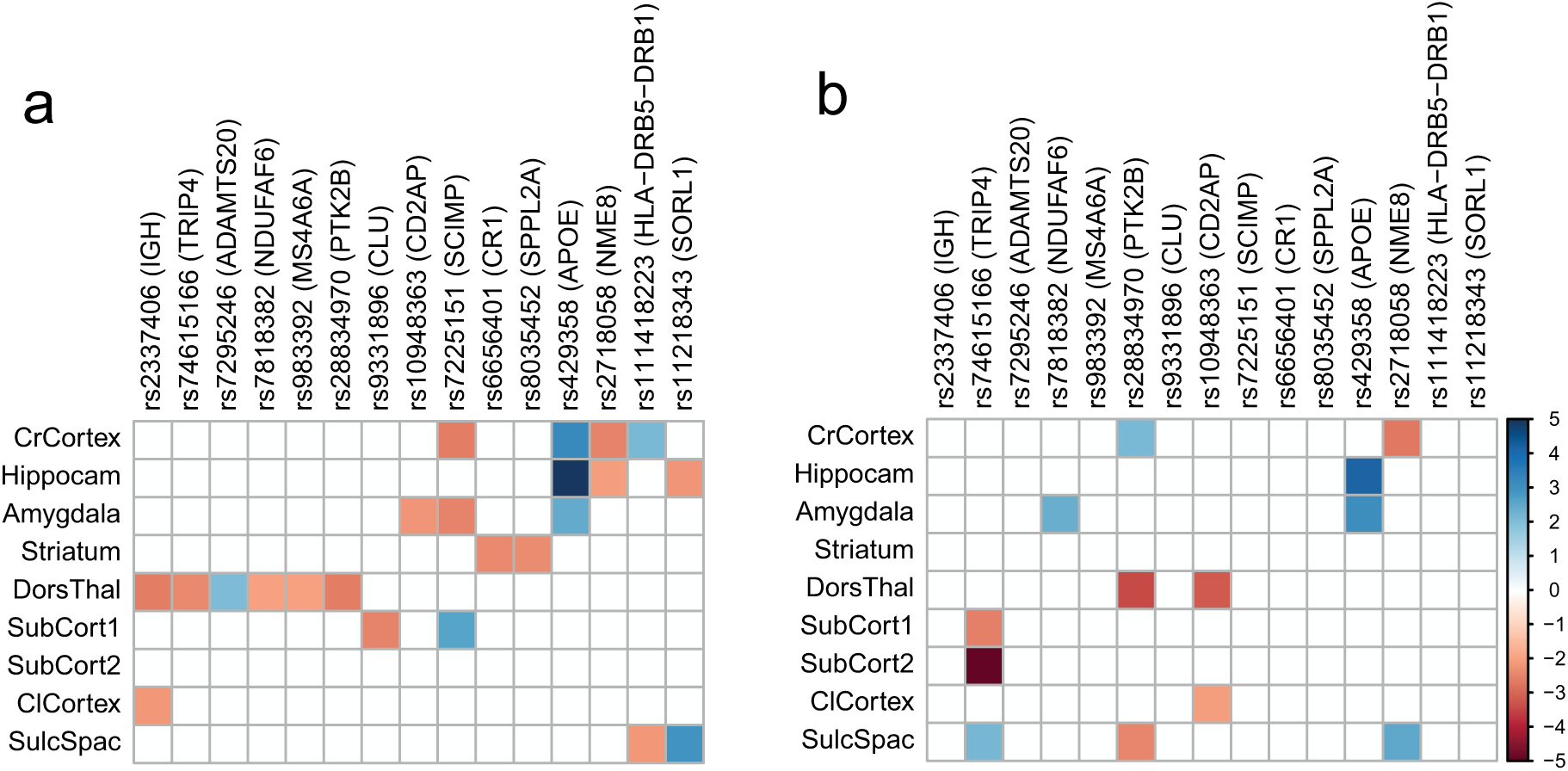
Association between SNPs and latent variable scores, as found by the robust maximum likelihood fit of the SEM. All nominally significant associations (*p* < 0:05) are coloured by their robust z-statistic values [20]. The linked genes [23] are shown in brackets. A: The results for the training set. After Bonferroni correction for the 315 test, only the effect of rs429358 (*APOE*) on the hippocampus region group remains significant. B: The validation results confirm the significant effect of rs429358 (*APOE*) on the hippocampus region group.

The latent variables reflect differences in brain region volumes across the ADNI dataset. To test whether these differences in brain region volumes were related to the disease phenotype, we compared the latent variable scores between the CN, LMCI, and AD individuals. Fig 5 shows the distribution of latent variable scores for the validation set. To calculate these, we used the fitted SEM of the training data, and used its parameter estimates to calculate latent variable scores for the validation data. For three region groups the latent variable scores were significantly lower in LMCI than in controls, and even lower in AD. These regions are the cerebral cortex, the hippocampal formation, and the amygdala. This reflects significant shrinkage in these areas during Alzheimer’s disease progression. The region group of sulci and spaces (SulcSpac) has a latent variable that significantly increases in LMCI and AD. The significant association between the SNP rs429358 and the latent variable scores for hippocampus reflects the importance of *APOE* for Alzheimer’s disease.

**Fig 5.**
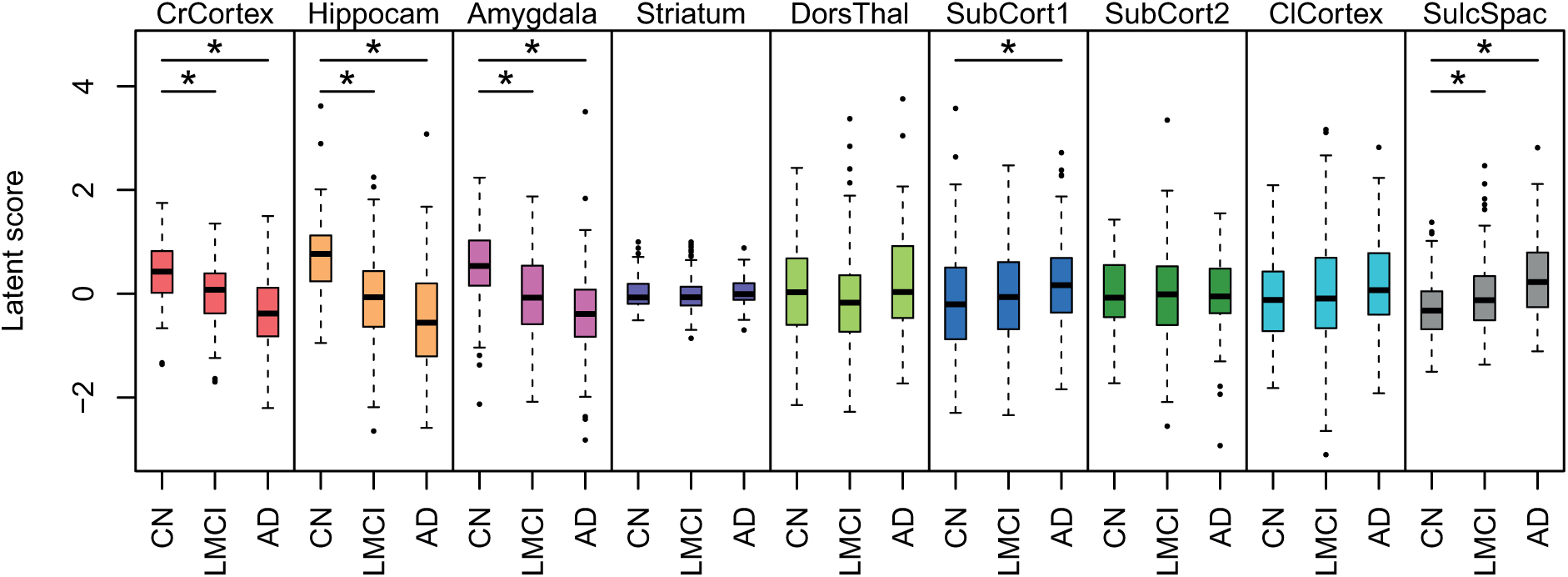
Association between the validation latent variable scores and diagnosis. Diagnosis is cognitive normal (CN), late mild cognitive impairment (LMCI), or Alzheimer’s disease (AD). Nominally significant differences (ANOVA *p* < 0:05) are indicated with asterisks. The cerebral cortex (CrCortex), hippocampus (Hippocam), and amygdala (Amygdala) latent volume variables are lowered with disease progression, while the latent variable score for sulci and spaces (SulcSpac) is increased.

## Conclusion

We have proposed the use of a maximum likelihood structural equation model for combining SNP data and structural brain area measurements. The model makes use of external gene expression data, to define groups of brain regions that may respond similarly to genetic variation. For each of these region groups, we define a latent variable, which captures the relationship between the regions in a group and genetic variation. We have applied the model on a simulated dataset, to show it can capture disease relevant variation and identify causal SNPs. In addition, we have applied the model to the ADNI dataset, containing Alzheimer’s patients, individuals with late mild cognitive impairment, and cognitive healthy controls. One SNP, linked to *APOE*, shows a reproducible significant relationship to the latent variable that captures hippocampal volume change. This latent variable, and that of the cerebral cortex, amygdala, and sulci & spaces also significantly associate with the disease diagnosis. This shows that our approach can be used to integrate several data types, and yield interpretable results.

The fitting process of the structural equation model has relatively high computational cost. It is truly multivariate, which makes it infeasible at the moment to perform genome-wide analysis. It does have advantages for incorporating a large number of variables, since it allows for straightforward inclusion of constraints on the parameter estimates [20]. With a constraint on the sum of squared weights, one could for instance implement a ridge regression. In addition, the model allows for the inclusion of additional data. This can be done either in the specification of the model structure, as we have done for the region groups, or by adding observed variables to the model.

These results show that maximum likelihood SEM is a versatile approach for data integration, which can be used to elucidate the relationships between genetic variation, structural brain phenotypes, and brain disease.

### Supporting information

**Supplemental Fig S1. Weights (loadings) from the latent variables to the region measurements.** The rows correspond to latent variables (region groups), and the columns to brain regions. Each first loading per region group was set to 1, for model identifiability. The loadings show the strength of the relationship between each latent variable and the thickness/volume of its corresponding brain regions. See Table S1 for the meaning of the region codes.

**Supplemental Table S1. All brain regions used in the ADNI section, with their region group code, manually annotated ABA region, ADNI code, and ADNI description.** Region group codes: *CrCortex* = cerebral cortex; *Hippocam* = hippocampal formation; *Amygdala* =amygdala; *Striatum* = striatum; *DorsThal* = dorsal thalamus; *SubCort1* =sub-cortical regions 1; *SubCort2* = sub-cortical regions 2; *ClCortex* = cerebellar cortex; *SulcSpac* = sulci and spaces.

## Acknowledgments

Data collection and sharing for this project was funded by the Alzheimer’s Disease Neuroimaging Initiative (ADNI) (National Institutes of Health Grant U01 AG024904) and DOD ADNI (Department of Defense award number W81XWH-12-2-0012). ADNI is funded by the National Institute on Aging, the National Institute of Biomedical Imaging and Bioengineering, and through generous contributions from the following: AbbVie, Alzheimer’s Association; Alzheimer’s Drug Discovery Foundation; Araclon Biotech; BioClinica, Inc.; Biogen; Bristol-Myers Squibb Company; CereSpir, Inc.; Cogstate; Eisai Inc.; Elan Pharmaceuticals, Inc.; Eli Lilly and Company; EuroImmun; F. Hoffmann-La Roche Ltd and its affiliated company Genentech, Inc.; Fujirebio; GE Healthcare; IXICO Ltd.; Janssen Alzheimer Immunotherapy Research & Development, LLC.; Johnson & Johnson Pharmaceutical Research & Development LLC.; Lumosity; Lundbeck; Merck & Co., Inc.; Meso Scale Diagnostics, LLC.; NeuroRx Research; Neurotrack Technologies; Novartis Pharmaceuticals Corporation; Pfizer Inc.; Piramal Imaging; Servier; Takeda Pharmaceutical Company; and Transition Therapeutics. The Canadian Institutes of Health Research is providing funds to support ADNI clinical sites in Canada. Private sector contributions are facilitated by the Foundation for the National Institutes of Health (www.fnih.org). The grantee organization is the Northern California Institute for Research and Education, and the study is coordinated by the Alzheimer’s Therapeutic Research Institute at the University of Southern California. ADNI data are disseminated by the Laboratory for Neuro Imaging at the University of Southern California.

